# Lyssa excreta: Defining parameters for fecal samples as a rabies virus surveillance method

**DOI:** 10.1101/2023.05.27.542586

**Authors:** Faith M. Walker, Jordyn R. Upton, Daryn Erickson, Zachary A. Barrand, Breezy Brock, Michael Valentine, Emma L. Federman, Emma M. Froehlich, Lolita Van Pelt, Lias Hastings, David L. Bergman, David M. Engelthaler, Crystal M. Hepp

**Affiliations:** School of Forestry, Northern Arizona University, Flagstaff, AZ; Pathogen & Microbiome Institute, Northern Arizona University, Flagstaff, AZ; TGen North Pathogen and Microbiome Division, Flagstaff, AZ; USDA APHIS Wildlife Services, Phoenix, AZ

**Keywords:** Lyssavirus, RABV, rabies, virus, Chiroptera, bat, non-invasive, disease surveillance, RT-PCR

## Abstract

It is not possible to systematically screen the environment for rabies virus (RABV) using current approaches. We sought to determine under what conditions RABV is detectable from feces and other accessible samples from infected wildlife to broaden the number of biological samples that could be used to test for RABV. We employed a recently-developed quantitative RT-PCR assay called the “LN34 panlyssavirus real-time RT-PCR assay”, which is highly sensitive and specific for all variants of RABV. We harvested and tested brain tissue, fecal, and/or mouth swab samples from 25 confirmed RABV positive bats of six species. To determine if rabies RNA lasts in feces sufficiently long post-defecation to use it as a surveillance tool, we tested fecal samples from 10 bats at the time of sample collection and after 24 hours of exposure to ambient conditions, with an additional test on six bats out to 72 hours. To assess whether we could pool fecal pellets and still detect a positive, we generated dilutions of known positives at 1:1, 1:10, 1:50, and 1:200. For six individuals for which matched brain, mouth swab, and fecal samples were tested, results were positive for 100%, 67%, and 67%, respectively. For the first time test to 24 hours, 63% of feces that were positive at time 0 were still positive after 24 hours, and 50% of samples at 72 hours were positive across all three replicates. Pooling tests revealed that fecal positives were detected at 1:10 dilution, but not at 1:50 or 1:200. Our preliminary results suggest that fecal samples hold promise for a rapid and non-invasive environmental screening system.

## INTRODUCTION

Rabies is a zoonotic disease of the central nervous system that invariably results in mortality [1]. It is caused by the RNA virus *Rabies lyssavirus* (RABV) and viruses from the *Lyssavirus* genus. RABV has the highest fatality rate of infectious diseases, with more than 59,000 human deaths globally each year [2]. Worldwide, dogs are the main RABV reservoir for human transmission, but in the Americas where vaccination of dogs is widespread, bats generate most of the human rabies cases [3, 4]. In Latin America, the primary species causing human infection is the common vampire bat (*Desmodus rotundus*) [5], while in the northwestern and southeastern U.S. tricolored (*Perimyotis subflavus*) and silver-haired (*Lasionycteris noctivagans*) bats have variants of rabies that are responsible for a higher proportion of human and terrestrial mammal deaths [6]. In Arizona, part of the American Southwest, suburban outbreaks of the disease occur regularly in wildlife populations, and interactions between wildlife and residents results in human exposures each year [7]. Arizona is one of the leading U.S. states for rabid wildlife, with bats, skunks, and gray fox the most common reservoir species [8]. Patyk et al. [9] found that among U.S. bat species, those in the Southwest were more likely to be rabid.

Despite its proximity and serious nature, it is not possible to systematically screen the environment for RABV. The gold standard for rabies diagnostics is the direct fluorescent antibody (DFA) test, which requires fresh brainstem tissues held at cold chain temperatures, requirements that prevent surveillance using inexpensive, field-collected samples [10]. Additional testing by the US Department of Agriculture, Animal and Plant Health Inspection Service, Wildlife Services is conducted through enhanced rabies surveillance using the direct rapid immunohistochemical test (DRIT) [11, 12] or as part of the rabies public health surveillance system. What is needed is a rabies detection approach based on readily available, non-invasive samples that can be applied broadly.

A recently-developed quantitative RT-PCR assay called the “LN34 panlyssavirus real-time RT-PCR assay” is highly sensitive and specific for all variants of rabies virus (RABV) [13, 14]. This assay was developed by a Centers for Disease Control (CDC) research group [14] and was found to be as or more sensitive than the DFA test [13]. It consists of a dual assay: the LN34 assay as well as a host species control β-actin real-time RT-PCR assay that signals presence of RNA in a sample and indicates PCR inhibition, extraction failure, or RNA degradation. Because the assay is highly sensitive, having succeeded with low quality and formalin-fixed samples, there is promise for non-traditional sample types such as feces, which contain intact and degraded nucleic acids [15]. Further, presence of RABV and other lyssaviruses in feces and saliva [16, 17] presents a surveillance opportunity with the LN34 assay, which has been shown to successfully detect RABV down to single digit copies of RNA [14].

Our overarching goal was to define and illustrate an effective and inexpensive surveillance system for rabies detection that can form the foundation of future statewide efforts to better understand public health risks. To address the goal, an environmental screening system for detection of RABV from feces first required evaluation of the strengths and limitations of feces as a potential sample type. We tested 1) RABV quantity in brain stem, mouth swab, and fecal material from infected individual bats; 2) evaluated RABV positivity of feces at ambient temperatures over 72 hours to better understand how long RABV may be detectable post defecation; and, 3) calculated RABV positivity of pooled fecal samples to determine how many fecal samples can be collected together to still return a positive result. We hypothesized that bat fecal samples could be reliably employed to detect RABV using the LN34 assay.

## METHODS

### Sample acquisition

This study was approved by the Institutional Animal Care and Use Committee (IACUC) at Northern Arizona University (Protocol 18-012). Carcasses and brain stems from bats found to be RABV positive via DRIT were provided by USDA Wildlife Services. Arizona bats evaluated included *Lasiurus xanthinus* (western yellow bat), *Eptesicus fuscus* (big brown bat), *Nyctinomops femorosaccus* (pocketed free-tailed bat), *Tadarida brasiliensis* (Mexican free-tailed bat), *Parastrellus hesperus* (western pipistrelle), *Lasiurus ega* (southern yellow bat), and *Antrozous pallidus* (pallid bat). Necropsies were performed in a BSL3 facility by staff with pre-exposure rabies vaccinations. We harvested feces from the intestines of bats using sterile scalpels and medical scissors, and used sterile cotton-tipped swabs to collect saliva. Samples were deposited into DNA/RNA Shield (Zymo Research, Irvine, CA, USA) and frozen at -80°C until RNA extraction.

### LN34 panlyssavirus real-time RT-PCR assay

We extracted RNA using the Zymo Direct-Zol RNA Miniprep Kit protocol. We performed the LN34 and β-actin RT-qPCRs on a QuantStudio 7 Flex (ThermoFisher Scientific, Waltham, MA, USA) as described previously [13, 14]. To summarize, for each sample, the LN34 assay targets the lyssavirus RNA genome and the β-actin assay targets host β-actin mRNA. Each 10 µL reaction contained Luna Probe One-Step RT-qPCR 4X Mix (New England Biolabs, Ipswich, MA, USA), primers (10 µM), probe (5 µM), and 2 µL RNA template. Samples were run as three replicates, and each assay run contained synthetic positive control RNA provided by the Center for Disease Control (Atlanta, GA, USA) and no template control reactions in triplicate. We used the LN34 assay diagnostic algorithm for post-mortem brainstem samples to determine the positive/inconclusive/negative thresholds [13].

### Tissue types, fecal time tests, and pooling

For individuals of five bat species (*Lasiurus xanthinus*, n = 1; *Lasiurus ega*, n = 1; *Nyctinomops femorosaccus*, n = 1, *Tadarida brasiliensis*, n = 1; *Parastrellus hesperus*, n = 2), we tested three tissue types (brainstem, saliva/mouth cells via mouth swab, and guano) with the LN34 assay. We also performed two time tests to determine how long feces remained positive at ambient conditions. We used feces of 1) ten RABV positive *Parastrellus hesperus* to 24 hours (0 and 24 hours), and, 2) six RABV positive bats (*Eptesicus fuscus*, n = 1; *Antrozous pallidus*, n = 1; *Lasiurus xanthinus*, n = 1;, *Tadarida brasiliensis*, n = 3) to 72 hours (0, 24, 48, and 72 hours). Each fecal sample was divided into two (for the 24 hour test) or four (for the 72 hour test) portions. When each time point was reached, we added 1 mL of DNA/RNA Shield to the fecal matter and stored the samples at -80°C until RNA extraction. To determine the extent to which fecal samples could be pooled in a field scenario (assuming the most dilute case of only one fecal sample from a RABV positive bat), we tested two known positive fecal samples (*Eptesicus fuscus* and *Tadarida brasiliensis*). We used 10 µL from 20 known RABV negative bat fecal extractions to make a negative pool. This was used to dilute the positive samples to 1:1, 1:10, 1:50, and 1:200.

## RESULTS

The six individuals for which we tested matched brain, mouth swab, and fecal samples, results were RABV positive for 100%, 67%, and 67% (Table 1), respectively. For the 24 hour time test, 63% of feces that were positive at time 0 were still positive after 24 hours (Table 2). For the 72 hour time test, all three replicates were positive for 50% of samples at 72 hours (Table 3). Pooling tests revealed that fecal RABV positives were detected at 1:10, but not at 1:50 or 1:200 (Table 4).

**Table 1.** Using the LN34 assay, rabies virus was detected in the brainstem, mouth swab, and feces of rabid bat carcasses. Values are Ct means of three replicates. LN=LN34, BA=β-actin.

| Bat species | Brainstem | Mouth swab | Feces |
| --- | --- | --- | --- |
| *Lasiurus xanthinus* | Positive (LN: 17.30, BA: 23.77) | Positive (LN: 25.15, BA: 31.58) | Positive (LN: 23.91, BA: 27.63) |
| *Nyctinomops femorosaccus* | Positive (LN: 22.25, BA: 25.10) | Inconclusive (LN: *, BA: 32.33) | Positive (LN: 30.24, BA: 23.57) |
| *Tadarida brasiliensis* | Positive (LN: 21.31, BA: 21.96) | Positive (LN: 31.45, BA: 30.98) | Inconclusive (LN: 40.16, BA: 28.52) |
| *Parastrellus hesperus* | Positive LN: (22.35, BA: 25.08) | Inconclusive (LN: *, BA: 33.41) | Positive (LN: 27.06, BA: 24.03) |
| *Lasiurus ega* | Positive (LN: 26.27, BA: 31.76) | Positive (LN: 30.19, BA: 37.14) | Inconclusive (LN: 38.87, BA: 36.99) |
| *Parastrellus hesperus* | Positive (LN: 19.32 BA: 27.04) | Positive (LN: 33.71, BA: 31.84) | Positive (LN: 31.09, BA: 31.50) |

\*Undetermined

**Table 2.** Positivity of fecal samples harvested from rabid *Parastrellus hesperus* bat carcasses and tested at time 0 and after 24 hours at ambient conditions, with mean Ct in parentheses.

|  | Time 0 | 24 hours |
| --- | --- | --- |
| 1 | Positive (LN: 24.51, BA: 25.25) | Positive (LN: 27.43, BA: 26.24) |
| 2 | Positive (LN: 31.98, BA: 30.51) | Positive (LN: 31.98, BA: 30.10) |
| 3 | Positive (LN: 32.00, BA: 30.10) | Positive (LN: 33.15, BA: 37.50) |
| 4 | Positive (LN: 30.61, BA: 25.15) | Positive (LN: 31.81, BA: 35.23) |
| 5 | Positive (LN: 24.89, BA: 28.80) | Positive (LN: 33.74, BA: 38.36) |
| 6 | Positive (LN: 33.61, BA: 28.12) | Inconclusive (LN: 35.54, BA: 35.40) |
| 7 | Inconclusive (LN: 37.18, BA: 29.55) | Inconclusive (LN: 37.56, BA: 29.22) |
| 8 | Inconclusive (LN: 37.01, BA 28.86) | Positive (LN: 32.51, BA: 34.46) |
| 9 | Positive (LN: 33.46, BA: 33.03) | Inconclusive (LN: 44.04, BA: 39.14) |
| 10 | Positive (LN: 32.46, BA: 31.36) | Inconclusive (LN: 36.82, BA: Undetermined) |

**Table 3.**
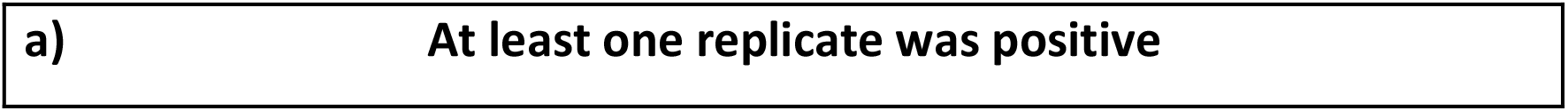
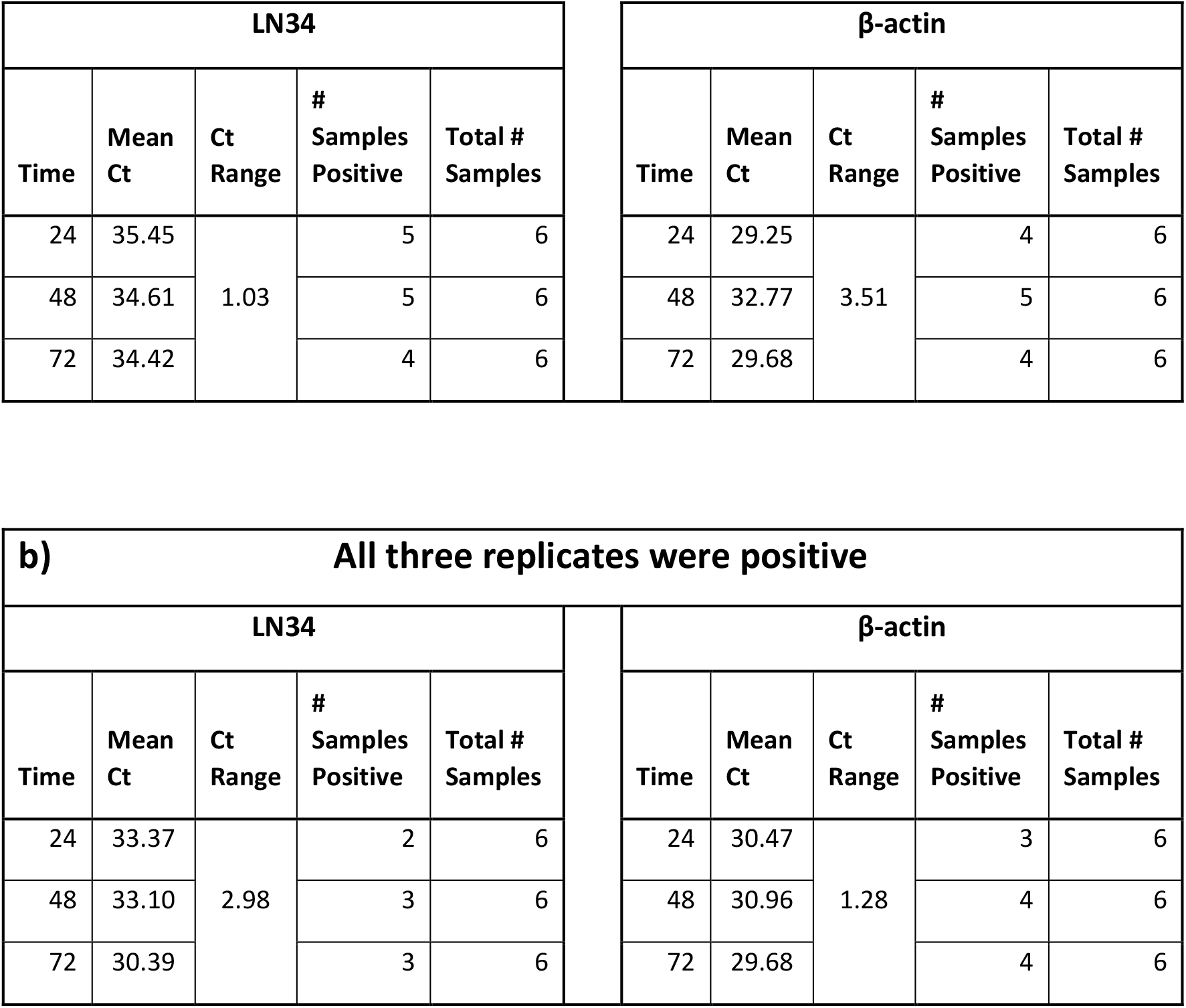
Positivity of fecal samples harvested from rabid bat carcasses and tested at time 0 and after 24, 48, and 72 hours at ambient conditions. At 72 hours, at least one replicate was positive for 4 of 6 samples (a) and all three replicates were positive for 3 of 6 samples (b). Bat species tested included *Eptesicus fuscus* (n = 1), *Tadarida brasiliensis* (n=3), *Antrozous pallidus* (n=1), and *Lasiurus xanthinus* (n=1).

**Table 4.** Positivity and amplification success of two RABV fecal samples declined at dilutions exceeding 1:10. Dilutions positive for rabies are <u>underlined</u>, LN is mean Ct over three replicates for the LN34 assay, the number of successful amplifications is in parentheses, and the positive control was a synthetic sequence provided by the Center for Disease Control. RABV fecal 1 was from a *Tadarida brasiliensis* and RABV fecal 2 was from an *Eptesicus fuscus*.

|  | 1:1 | 1:10 | 1:50 | 1:200 |
| --- | --- | --- | --- | --- |
| **RABV fecal 1** | LN: <u>31.25</u>, BA: 26.52 (3) | LN: <u>34.12</u>, BA: 29.73 (3) | LN: 36.56, BA: 30.56 (1) | Undetermined (0) |
| **RABV fecal 2** | LN: <u>32.13</u>, BA: 35.71 (3) | LN: <u>34.82</u>, BA: 31.93 (3) | LN: 35.56, BA: 32.17 (1) | Undetermined (0) |
| **Positive control** | LN: <u>27.65</u>, BA: undetermined (3) | LN: <u>30.46</u>, BA: 31.8 (3) | LN: <u>32.32</u>, BA: 31.81 (3) | LN: <u>34.14</u>, BA: 31.84 (3) |

## DISCUSSION

Bat fecal samples hold promise as a surveillance method for rabies virus. The successful testing of feces, for which nucleic acids are naturally degraded and in a matrix containing multiple inhibitors [15], was likely aided by a short amplicon; the LN34 assay amplifies a 165 bp region. We found that fecal samples of postmortem rabid bats yielded PCR Ct values that were higher than that of brain tissue (*i*.*e*., lower quantity), but the majority remained positive. Half of the fecal samples were positive to at least 72 hours at ambient temperatures, which suggests that there is time post-defecation to collect feces and for rabies to still be detectable. Feces could still be detected when pooled at a ratio of 1:10 (one fecal pellet with rabies virus collected together with nine without rabies virus), which provides guidance for pooling of feces in the field. Notably, we used the established Ct thresholds for brain tissue, but it may be that new, higher thresholds to assign positivity could be set for this sample type [13].

Fecal samples will allow determination of rabies presence at a site or region, and will do so at a scale that is not currently possible with postmortem tissues. It will be possible, for example, to non-destructively sample bat roosts to determine the enzootic prevalence and seasonality of RABV. The assay is inexpensive, and thus it is possible to sample broadly, and to do so in any locale that has RT-PCR capability. Further, it is likely that feces from mammals other than bats can be targeted. For instance, surveillance programs using canine feces would benefit vaccinations campaigns and the effort to eliminate dog-mediated human rabies deaths by 2030 [18]. Finally, it will be possible to explore pairing positive fecal samples with sequencing methods to determine the phylogeographic dynamics of species specific variants and better understand the evolving risk of zoonotic expansion.

